# Computational Structure Prediction Provides a Plausible Mechanism for Electron Transfer by the Outer Membrane Protein Cyc2 from *Acidithiobacillus ferrooxidans*

**DOI:** 10.1101/2021.03.22.436458

**Authors:** Virginia Jiang, Sagar D. Khare, Scott Banta

## Abstract

Cyc2 is the key protein in the outer membrane of *Acidithiobacillus ferrooxidans* that mediates electron transfer between extracellular inorganic iron and the intracellular central metabolism. This cytochrome c is specific for iron and interacts with periplasmic proteins to complete a reversible electron transport chain. A structure of Cyc2 has not yet been characterized experimentally. Here we describe a structural model of Cyc2, and associated proteins, to highlight a plausible mechanism for the ferrous iron electron transfer chain. A comparative modeling protocol specific for trans membrane beta barrel (TMBB) proteins in acidophilic conditions (pH ~2) was applied to the primary sequence of Cyc2. The proposed structure has three main regimes: extracellular loops exposed to low-pH conditions, a TMBB, and a N-terminal cytochrome-like region within the periplasmic space. The Cyc2 model was further refined by identifying likely iron and heme docking sites. This represents the first computational model of Cyc2 that accounts for the membrane microenvironment and the acidity in the extracellular matrix. This approach can be used to model other TMBBs which can be critical for chemolithotrophic microbial growth.

**Importance of work:** *Acidithiobacillus ferrooxidans* can oxidize both iron and reduced sulfur compounds and plays a key role in metal sulfide ore bioleaching used for the industrial recovery of metals. *A. ferrooxidans* has also been explored as a potential organism for emerging technologies such as e-waste recycling and biofuel production. Synthetic biology efforts are hampered by lack of knowledge about the mechanisms of iron oxidation and reduction, which is mediated by the Cyc2 transmembrane beta barrel (TMBB) protein.

## Introduction

*Acidithiobacillus ferrooxidans* is a gram-negative acidophilic chemolithoautotroph that exhibits the highest growth rates in the range between pH 1.8 and 2.2 (37). It is an important electrochemically active bacterium (EAB) involved in mineral bioleaching due to its ability to oxidize both ferrous iron and reduced sulfur compounds. This capability has been leveraged in industrial processes where the resulting oxidized iron is exploited for the recovery of copper and other valuable metals from sulfidic ores (67; 85) (Fig. 1a). In addition to its role in industrial “biomining” there has been increasing interest in developing *A. ferrooxidans* for other applications such as electronic waste recycling (72), and as a unique chassis organism for biofuel production. Genetically modified *A. ferrooxidans* cells have been engineered to create biochemicals from CO_2_ using electrochemically reduced iron as the sole energy source, and these types of “electrofuels” platform provide a promising route towards enabling the conversion of renewable electrical energy into the chemical energy of transportation fuels (51; 42; 10).

**Figure 1.**
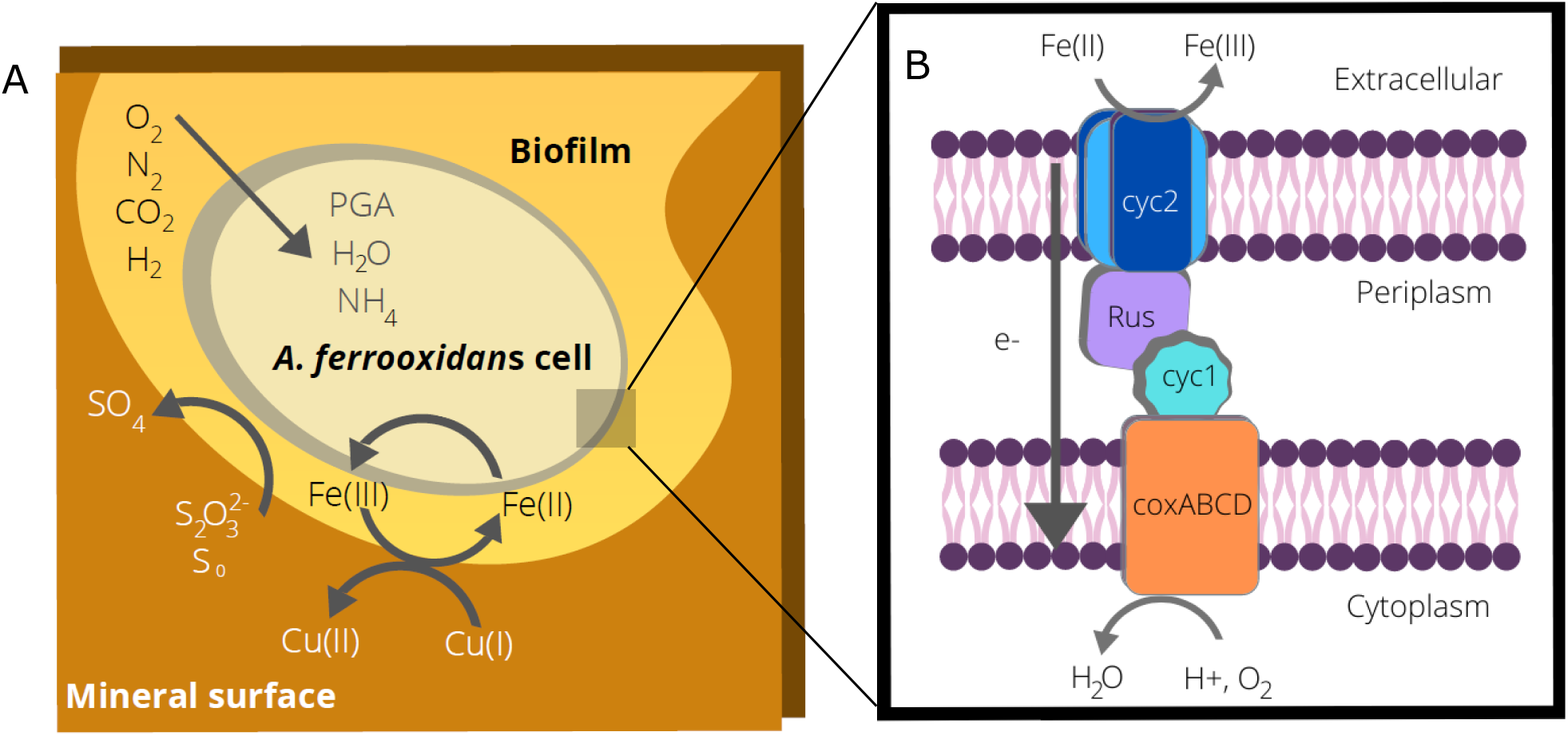
Overview of metal oxidation by *A. ferrooxidans*. A. Overview of *A. ferrooxidans* metabolism when growing on ores containing sulfide minerals such as chalcopyrite (CuFeS_2_). The bacterium oxidizes iron and reduced sulfur compounds, resulting in the solubilization of copper or other commercially valuable metal targets. Carbon dioxide is fixed via the Calvin cycle to 3-phosphoglycerate (PGA). B. Diagram of the electron transfer pathway used for extracellular iron oxidation. Electrons are transferred from the extracellular matrix, through the outer membrane (Cyc2), the periplasmic space (Rus and Cyc1) to coxABCD in the inner membrane. This is a simplified model of the downhill electron transfer pathway, as other periplasmic proteins may also be involved in this process. Under some conditions, electron transfer can be reversed, and iron reduction can also occur.

When not growing on sulfur, *A. ferrooxidans* cells are able to obtain all of their metabolic energy via the extracellular oxidation of ferrous iron. Thus iron supports mediated electron transfer (MET), enabling the cells to grow planktonically and the iron can be reduced by solid surfaces or by electrodes. The cells can also demonstrate direct electron transfer (DET) where they are able to grown on solid surfaces including electrodes, and the electrons can flow directly into the cellular metabolism (18). Once inside the cells, the electrons flow through two pathways. In the thermodynamically uphill pathway, electrons are used to regenerate the reducing equivalents (NADP(H)). In the thermodynamically downhill pathway, oxygen is the final electron acceptor and protons are used to produce water, enabling the cells to produce ATP and to maintain an internally neutral pH (77). When growing on reduced sulfur compounds, oxygen usually serves as the terminal electron acceptor. However, under anaerobic conditions, ferric iron can also serve as an electron acceptor (73; 48). Therefore, under these conditions, the flow of electrons is reversed and the cells become iron reducers. Thus, unlike many other EAB, the electron transfer machinery in *A. ferrooxidans* is reversible.

The proteins involved in electron transfer into and out of the cells are encoded in the rus operon; which includes an outer membrane cytochrome c protein (Cyc2), a periplasmic blue copper protein (rusticyanin, or Rcy), a periplasmic cytochrome c4 (Cyc1), and an inner membrane aa3-type cytochrome c oxidases (7; 99) (Fig. 1b). The periplasmic proteins Rcy and Cyc1 have previously been crystallized and characterized (87; 1), and the surface interactions at the interface between Rcy and Cyc1 have been determined (66). The Cyc2 protein (485 amino acids, MW 52.4 kDa, Fig. S1, Fig. S2) is the critical outer transmembrane protein that serves as the extracellular interface for electron transfer necessary for iron oxidation or reduction (97; 18). Overexpression of the *cyc2* gene increases Fe(II) oxidation (54), as does overexpression of the *rus* gene that encodes Rcy (36). 3-D structural information for Cyc2 has not yet been reported, likely due to experimental difficulties, and this hampered our mechanistic understanding of this key energetic pathway.

Computational modeling of the Cyc2 protein structure has been difficult as there are few homologous membrane protein structures available based on sequence similarity that have been crystalized. The development of robust computational methodologies that can account for the membrane microenvironment is a growing field of research (2; 69). Outer membrane proteins in gram-negative bacteria typically contain transmembrane beta barrel (TMBB) domains that act as pores to control intercellular transport (15). As Cyc2 is located in the outer membrane, it is likely a TMBB. To date, most transmembrane protein modeling efforts have focused on alpha-helical transmembrane proteins, which are most often found in the inner cell membrane (44; 12; 13; 89); however modeling and designing TMBBs is an emerging area of research. The 3D structural prediction of TMBBs from primary amino acid sequence is hindered by limited structural information, however several tools have been developed for both comparative (CM) and *de novo* modeling of TMBBs (75; 30; 3; 32; 27; 81). Computational tools for *de novo* design of TMBBs have also been developed (79; 86). Additionally, as *A. ferrooxidans* require highly acidic environments, modeling methods applied to Cyc2 should account for pH-dependent protein conformations.

Experimental evidence indicates that Cyc2 has several cofactors and binding interactions. Ferrous iron does not accumulate significantly within the cells, and the ferrous oxidation site is localized in the outer membrane facing the exterior of the cell (38). Cyc2 domains have been experimentally demonstrated to project from the cells into the extracellular matrix (97). The N-terminus of the protein primary sequence contains one *c*-type cytochrome motif CXXCH, where a heme likely is covalently bonded to the protein by two thioether bonds with the cysteine sulfur atoms and an H-bond to the nitrogen on the histidine ring (4). Purified Cyc2 shows a monohemic signature when characterized by spectrophotometry (97).

To create structural models of Cyc2 and related outer membrane proteins, there is a need to develop TMBB-specific modeling tools so that new insights can be gained into how electrons are transferred from external sources to interior cellular metabolisms. Rosetta, a widely used biomolecular modeling and design package, contains computational frameworks for comparative modeling (RosettaCM)(13), membrane protein design (RosettaMP)(2), and pKa calculations (Rosetta-pH)(43). Additionally, there are existing Rosetta protocols for determining protein-protein docking interactions (Rosetta Dock). RosettaCM has a well-documented protocol for comparative modeling of alpha helical transmembrane proteins but no specialized protocol for TMBB structures currently exists (25).

In this manuscript, these existing Rosetta function platforms were combined to develop a TMBB model for the Cyc2 cytochrome c protein of *A. ferrooxidans*. The modeling framework was adjusted for the environmental conditions specific for *A. ferrooxidans.* Putative metal binding sites in the structure were identified, and docking of the Rus and Cyc1 proteins was explored. This new structural ensemble can be used to provide insights into the formation of the natural “molecular wire” that enables reversible electron exchange with inorganic materials in the environment.

## Results

The Rosetta membrane protein comparative modeling protocol has been primarily used to model alpha helical transmembrane proteins (96). The protocol involves a low-resolution step where amino acid sidechains are represented by a single pseudo-atom placed at the geometric centroid of all atom positions (centroid mode) followed by a high-resolution step in which all atoms including hydrogen atoms are explicitly modeled (full-atom mode). Interactions in the low- and high-resolution steps are described using the centroid and full atom energy functions, respectively. To expand the scope of this methodology to include TMBB proteins like Cyc2, we altered the low-resolution centroid score function by removing alpha helical membrane protein-specific terms from the low-resolution centroid score function (*mp_nonhelix, mp_termini, mp_tmproj*) (Fig. S3). Next, the full atom score function was modified to *franklin2019* weights, which includes fa_water_to_bilayer, a score term that accounts for membrane-embedded protein transfer free energy (2). We tested the impact of these changes on TMBB structure prediction on three test proteins as described in the Methods/SI and found significant improvements in prediction performance (Fig. S4, S5).

With a protocol for modeling TMBB proteins in hand, we turned to the Cyc2 primary protein sequence from *Acidithiobacillus ferrooxidans* ATCC 23270 (Fig. S1). The modified RosettaCM protocol for TMBB proteins was used to model Cyc2 from PDB IDs 2ov4 and 4rjw (65; 80), homologous 16-stranded TMBB proteins in the OPM database (Fig. 2a). As the outer membrane of *A. ferrooxidans* faces an extracellular matrix at pH 2, the full atom score function was modified to include terms *fa_elec* and *e_pH* and the Rosetta-pH package was initialized to match known pH values for the extracellular matrix. Salt bridges between side chains that were present at pH 7 were not predicted at pH 2 (Fig. S6), which is consistent with experimentally observed salt bridge formation behavior (64). The lowest-scoring model was scored by MolProbity where it had an overall MolProbity score of 2.01, falling into the 73^rd^ percentile of all available structures based on assessments of protein geometry. Ramachandran favored rotamers comprised 98.62% of the residues, while Ramachandran poor rotamers comprised 0.27% of the protein. The Cyc2 model had a ProQ3 S-score of 0.36, which is within the predicted range of scores for proteins in the CASP11 test set (0.40 ± 0.12) (83; 82). The sequence for Cyc2 was accessed in PSIPRED for secondary structure and domain topology (Fig. S7) (57; 17).

**Figure 2.**
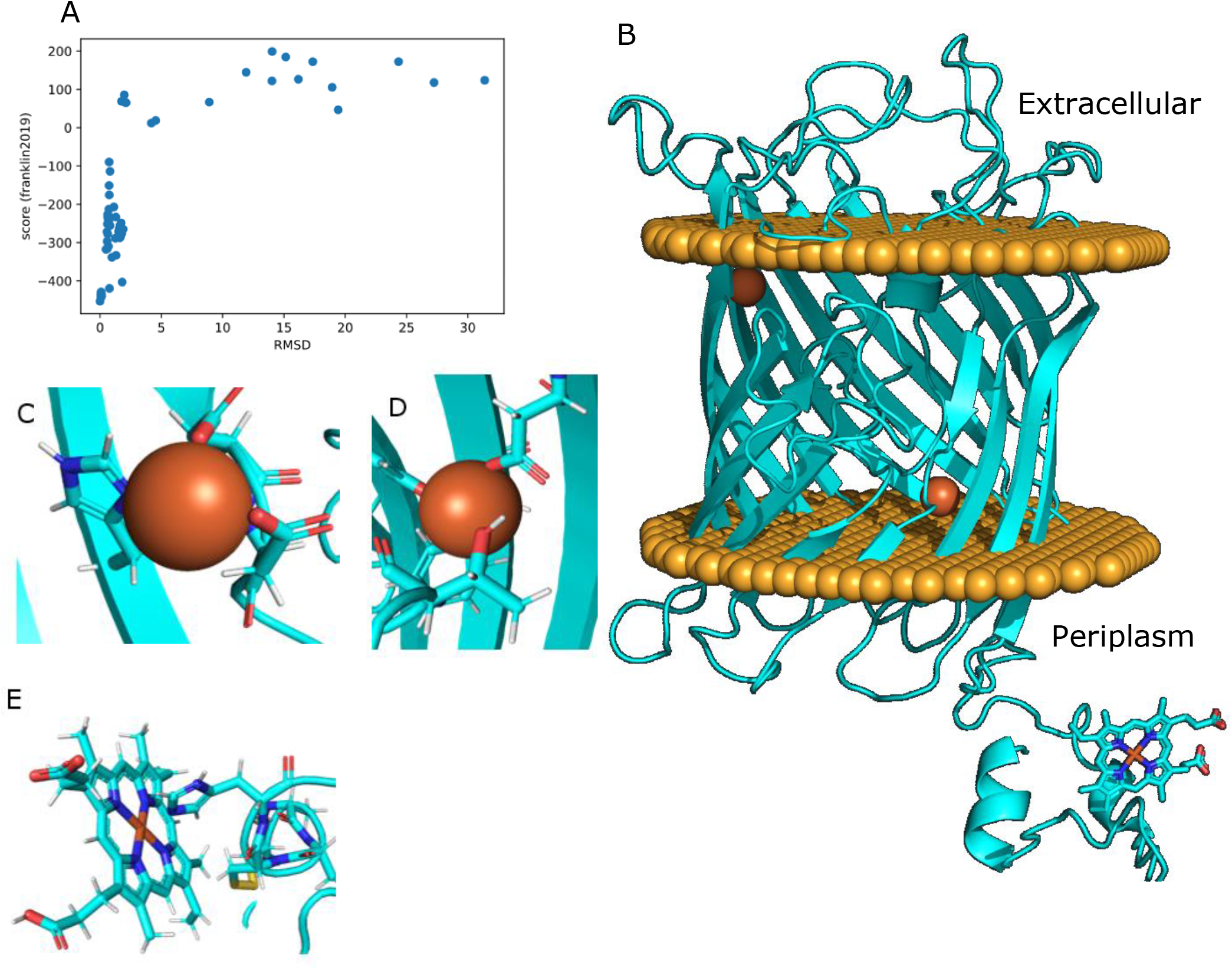
Characterization of modeled Cyc2 protein and interactions with ligands: A: Funnel plot for one hundred Cyc2 modeling trajectories using the modified set of membrane weights. The c-alpha RMSD was measured in reference to the lowest-scoring structure. B: Lowest-scoring Cyc2 protein model. Disks of spheres represent the phospholipid heads on the outer membrane where the top of the protein protrudes into the extracellular matrix and bottom of the protein (including the heme group) is in the periplasm. C: Square planar iron chelating geometry for residues H119, D137, D138. D: Tetrahedral iron binding geometry for Y262 and D308. E. Binding interaction between H16 and heme c.

The resulting model is a monomeric 16-stranded TMBB that spans the membrane, with a cytochrome-like domain consisting of several short alpha helices and a bound heme group that is fused to the beta barrel at the N-terminus of the protein (Fig. 2b). The TMBB topology is similar to other 16-stranded TMBB porins. The model has an annular pore at the apical side facing the periplasmic space, with flexible loops facing the acidic extracellular environment and the N-terminus cytochrome-like domain protruding into the periplasm. The majority of the protein contains beta-sheet rich domains, as expected for an outer membrane TMBB protein.

Potential metal binding sites were then explored in the Cyc2 model structure. Homologous binding locations were identified against all known metal binding structures in the RCSB PDB using the Metal Ion-Binding Site Prediction and Docking Server (MIB) (Fig. S8). The binding sites with evolutionarily conserved residues as measured by ClustalOmega in ten related iron-oxidizing bacteria were identified and used as initial binding sites for metal ion docking (60) and metal chelation geometry was optimized using Rosetta (Fig. S9). Two binding sites were identified. The most likely binding location was at residues H119, D137, D138, and was homologous to a ribonucleotide reductase protein from *Corynebacterium ammoniagenes* (PDB ID: 1kgo)(34). Calculated binding geometries are square planar, with angle RMSDs of 6.2° (Fig. 2c). An additional probable second binding site was identified at Y262 and D308 and was homologous to a site in YmdB, a phosphodiesterase from *Bacillus subtilis* (PDB ID: 4b2o)(22). The binding geometry of this site is tetrahedral, with angle RMSDs of 7.3° (Fig. 2d).

To further investigate electron transfer pathways in the predicted Cyc2 structure, the model was further refined by docking the heme C binding domain via homology to a heme binding protein involved in the electron transfer pathway in *Vibrio parahaemolyticus* by aligning the CXXCH motifs (PDB ID: 2zzs)(14). Calculated binding geometries are octahedral, with angle RMSDs of 2.7° from ideal (Fig. 2e).

The predicted ligand and ion binding interactions with the model Cyc2 structure are consistent with prior experimental work in *A. ferrooxidans.* The structure has two internally bound iron atoms in the barrel region and one bound heme c in the cytochrome-like region (Fig. 2b), which corresponds with ligands that have been co-eluted with Cyc2 during purification (6). Heme-protein interactions fall within experimentally measured parameters. The imidazole ring of H16 is 2.1 A from the docked central iron in the heme C (Fig. 2e), which corresponds to the experimentally determined interaction radius (68; 101). Metal residue chelating geometries around the bound irons have low angle RMSD values below 10°, indicating that structural geometries of the coordination spheres approach the ideal metal ion environments within the protein (95).

Next, docking was performed to find possible interfaces between Cyc2 and its downstream partners in the electron transfer chain, Rcy and Cyc1. These are periplasmic proteins with crystallographic structures available (PDB ID: 1rcy and 1h1o) (87; 1). A structural model of the Rcy-Cyc1 complex has been previously predicted: Abergel et. al determined a probable interface between of rusticyanin and Cyc1 (Fig. S10) We confirmed this interface was plausible by using ClusPro for FFT patch-based docking and refining using local Rosetta docking. The modeled Rcy-Cyc1 complex was then docked with Cyc2 to produce an interface score vs RMSD plot with a narrow energy funnel. Patches were identified by regions of conservation with electron transfer-active or chelating residues on Rcy. D58 was identified as a probable patch location on Rcy, with H57 as a possible redox center. A similar hydrogen bonding interface was identified on the docked model of Cyc2 to Rcy (Fig. S10).

The protein-protein docking results are consistent with experimental findings. Docking simulations with Rosetta confirmed previous modeling efforts to dock Rcy and Cyc1. H57 and D58 in Rcy had previously been thought to be a possible copper binding site prior to crystallographic analysis (78), as the residues on Rcy in those positions align with known copper binding sites on azurin from *Pseudomonas aeruginosa*. By contrasting extended X-ray absorption fine structure (EXAFS) on oxidized and reduced forms of Rcy, it has been shown that there are multiple rigid electron transfer sites potentially contributing to the higher reduction potential of Rcy (35). The authors identified electron transfer interactions involving nitrogen atoms on histidine imidazole rings (H57, H85, and H143), which had all previously been thought to be involved in copper coordination (28). However, the X-ray crystal structure of Rcy showed that two of the identified histidines coordinate copper binding (H85 and H143). Large interatomic distances between H57 and the copper ion make it unlikely that all the histidines chelate the same copper ion. We hypothesize that H57 could instead play a role in the electron transfer pathway between Cyc2 and Rcy, and D58 may be involved in stabilizing the interface between Cyc2 and Rcy due to its hydrogen bonding ability (66). Therefore, this residue patch is thought to be most likely Cyc2-Rcy interface on Rcy.

The docked structures were used to identify putative electron transfer pathways through the full Cyc2-Rcy-Cyc1 complex (Fig. 3a). Emap, a platform that identifies potential electron tunneling pathways via using graph theory to map node distances between aromatic amino acids, was used to discover potential transfer pathways (80). This analysis indicates there exists a probable electron hopping pathway within the full model complex from the Cyc2 model structure to the experimentally verified Rcy-Cyc1 complex (Fig. 3b). The residues implicated in electron transfer with this model of Cyc2 are primarily phenylalanines and tyrosines, a phenomenon which has been experimentally observed in yeast cytochrome c via point mutations to residues without electron-transfer active moieties (52). The computational predictions for Cyc2 account for known electron transfer interactions, such as electron hopping between Rcy H143 and the Rcy docked copper ion. Additionally, there are experimentally predicted electron interactions in Cyc1 between Y143, Y42, and the most downstream bound heme (heme 1185) (1). The shielding provided by the tyrosine residues around Cyc1 heme 1185 indicates that heme 1185 is less solvent-accessible than heme 1184 and therefore heme 1185 is the higher potential driving force in the Cyc1 cytochrome.

**Figure 3.**
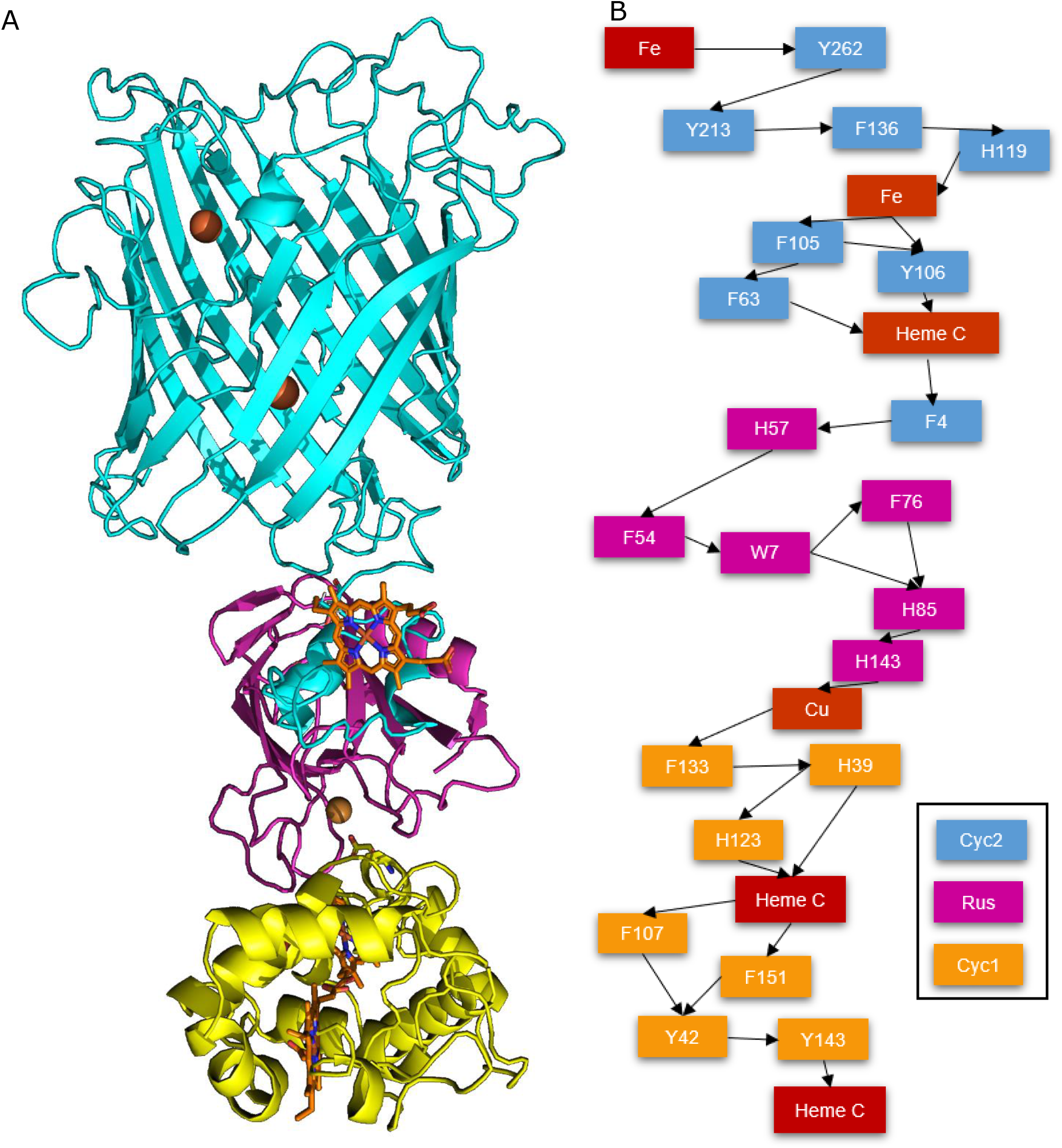
Predicted protein-protein interactions supporting the electron transport chain: A: Docked models of the Cyc2-rusticyanin-Cyc1 pathway (where carbon backbone of Cyc2 is blue, rusticyanin in magenta, and Cyc1 is in yellow). Hemes, iron ions, and copper ions are indicated in sepia. B: Predicted electron hopping pathway for electron transfer from the exterior of the cell into the periplasm of *A. ferrooxidans*.

Possible Fe(II) transient binding sites were identified using regions of negative potential as calculated by Poisson-Boltzmann scores (Fig. 4a, 4b). Within this model, a putative iron binding site on the exterior of the cell may be present, with a topological pocket in a region of high negative potential. However, more experimental evidence will be needed to verify that this is the site of iron binding in Cyc2. When viewed from the periplasmic side of the protein, the solvent-exposed pore extending through the protein has highly negative Poisson-Boltzmann potential (Fig. 4c). This is consistent with porins selective for cations; porins selective for anions have inner pores with positive Poisson-Boltzmann potentials, while porins with no ion selectivity show no clear pattern in Poisson-Boltzmann potential (Fig. S11). Thus the negative potential of the pore suggests Cyc2 may also facilitate cation transport.

**Figure 4.**
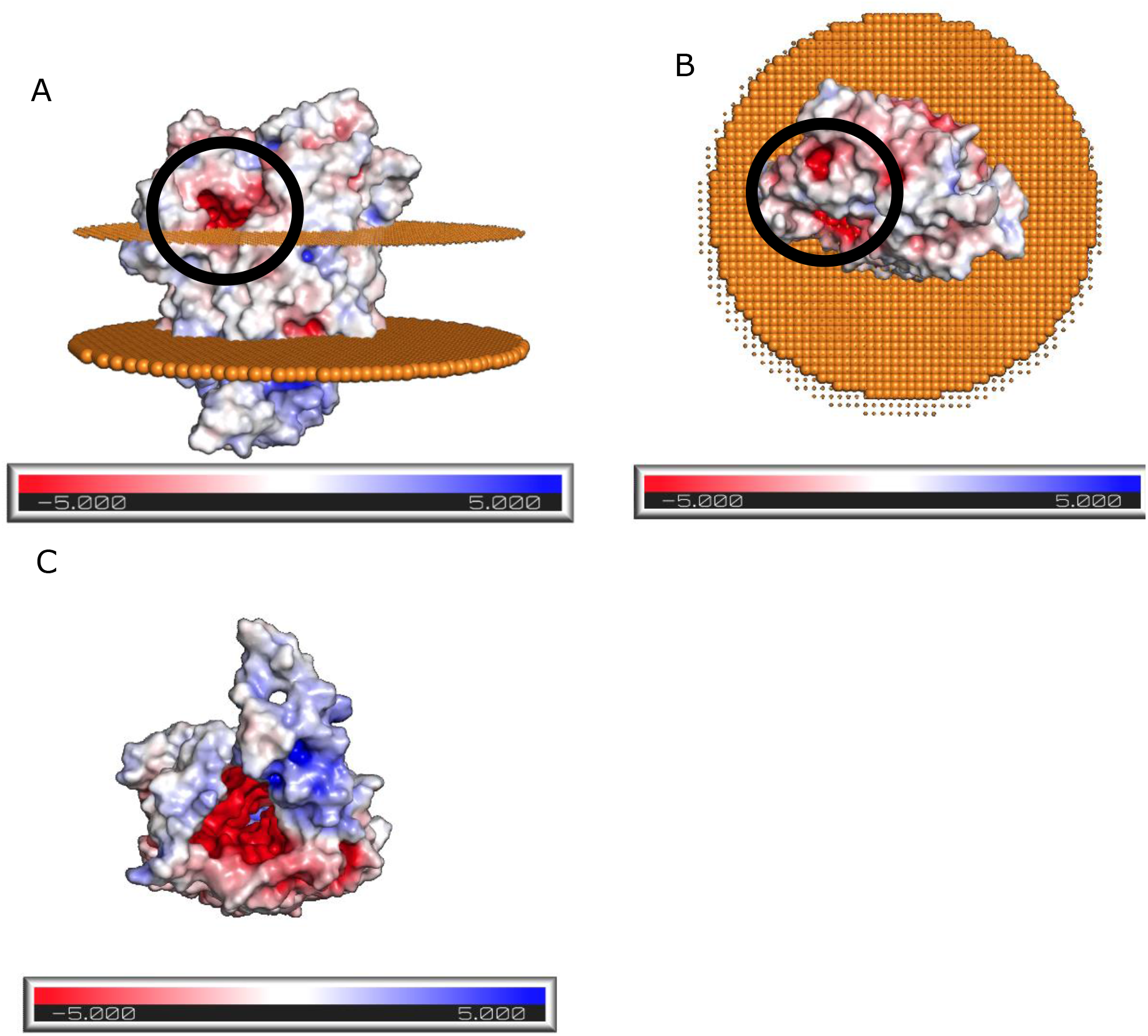
Poisson-Boltzmann potential identifies possible iron binding pocket on the extracellular region of Cyc2: Poisson-Boltzmann potential for Cyc2, where blue indicates regions of positive potential (> +5 kT/e) whereas red depicts negative potential (< −5 kT/e). A: labeled potential viewed from the side, where the upper disk represents phospholipid heads on the outer membrane facing the extracellular matrix and bottom disk represents the phospholipids facing the periplasm. There exists a region of high negative potential (circled in black) that may serve as a possible iron binding site. B: Labeled potential as viewed from above, showing the protein structure protruding from the cell. Regions of high negative potential are circled in black. C: Labeled potential as viewed from below the membrane, as would be observed from inside the cell. An annular space exists within the protein, inside which is predicted to have a high negative potential

## Discussion

Cyc2 is the key outer membrane protein in *A. ferrooxidans* that enables electron transfer from extracellular inorganic materials (19; 49; 47). To create a structural model, a modified score function was developed which allows Rosetta comparative modeling protocols to be extended to TMBBs. This procedure was used to produce a new structural model of the Cyc2 protein structure. The heme ligand was docked onto the protein and putative metal binding sites were identified. The modelled protein was docked onto Rcy-Cyc1, leading to a model of the electron transfer pathway in *A. ferrooxidans* that is consistent with known experimental details.

EAB all share the ability to exchange electrons across their insulating outer membrane(s), although different mechanisms have evolved depending on the specific metabolism of the species. In every case, electron transfer is mediated by cofactor-containing outer membrane proteins (91; 93). For example, *Shewanella oneidensis* is a metal oxide-reducing gram-negative bacterium, which uses a combination of methods for electron uptake but is dominated by flavin-dependent MET (45; 70; 94). The structure of the main *S. oneidensis* transmembrane electron conduit MtrCAB is trimeric: one peptide protrudes into the extracellular matrix, one serves as a transmembrane porin, and one is a multi-heme cytochrome (26). There is a directional hierarchy in the electron transfer pathway due to the presence of multiple chained hemes (29).

Another species of EAB, *Geobacter sulfurreducens,* transfers electrons from mineral surfaces using conductive pili (76; 61). Over 110 genes coding for putative c-type cytochromes have been identified in *G. sulfurreducens,* resulting in many parallel electron transfer pathways (63). Two major trimeric conduits have been identified, OmaB-OmbB-OmcB and OmaC-OmbC-OmcC, both of which consist of two cytochromes and one transmembrane porin (56; 55). *G. sulfurreducens* electron transfer can be bidirectional, but experimental findings suggest the cathodic electron transfer, where electrons move from interior metabolism to the exterior matrix, may take place via an alternative non-heme protein (33).

*G. sulfurreducens* and *S. oneidensis* have the most extensively studied porin-cytochrome systems which are both trimeric systems. There are also some dimeric porin-cytochrome systems, including *Sideroxydans lithotrophicus,* and *Rhodopseudomonas palustris. A. ferrooxidans* lacks a multimeric porin-cytochrome complex; rather, Cyc2 is a simpler monomeric fused porin-cytochrome structure that enables electrons to flow in or out of the cell depending on the orientation of the driving force (91).

This *A. ferrooxidans* Cyc2 structure may be useful for developing structures for homologous TMBB electron transfer complexes. For instance Cyt572, a functionally and structurally similar protein with low sequence identity to Cyc2, has been identified in other acidophilic iron-oxidizing bacterial communities comprised of *Leptospirillum* spp (39). And the *cyc2* gene is conserved across a range of neutrophilic iron-oxidizing bacterial mats (11; 62; 41). Currently, there are no crystallized structures of functionally homologous fused porin-cytochromes found in the outer membrane in the OPM databank (58).

It has been suggested that the electron transfer from Fe(II) to Cyc2 occurs via a transient encounter-Michaelis complex (16; 50), such that there is a transient iron-binding site on the outside face of Cyc2 that serves as the initial binding site for Fe(II) oxidation and electron transfer. Such transient interactions are not well-captured by crystallography, so there are no homologous binding geometries in the Protein Data Bank. As shown in Fig 4, there is a putative metal binding site on the Cyc2 protein structure that is close to the outer membrane surface. An additional poorly understood feature of Cyc2 is the binding site specificity between different metal ions. In addition to growth on Fe(II) it has also been shown that the cells can oxidize uranium from uranous to uranyl, suggesting that the Cyc2 electron transport chain can carry electrons from both uranium and iron (23; 90). It has also been demonstrated that the cells cannot grow on vanadium (III). However when a small amount of iron is included, growth on V(III) can occur where iron is used as a redox mediator between vanadium and the cells (51). The mechanism for this metal selectivity is not yet known and further experimental work will be necessary to validate the location of the metal binding pocket.

## Conclusion

A comparative modeling protocol was applied to the Cyc2 sequence to obtain a modular model with a cytochrome-like heme c binding domain, a flexible linker, and a membrane-embedded beta barrel. Iron-binding sites within Cyc2 were identified by homology. Protein-protein docking was performed to determine likely interfaces between Cyc2, Rcy, and Cyc1 to produce an “conducting wire” spanning from the extracellular matrix through the outer membrane into the periplasm. Electron hopping distances in the pathway were calculated using graph theory, and it was determined that there is a plausible electron transfer pathway from Fe(II) through the model of Cyc2 and to the docked Rcy-Cyc1 complex. This model is the only known computational model of Cyc2 that accounts for the membrane microenvironment and the acidity levels in the extracellular matrix and aligns with experimental findings. This model of Cyc2 provides a blueprint for the genetic engineering of Cyc2. And the modeling approach may be useful for modeling other TMBBs which can be critical for chemolithotrophic microbial growth.

## Materials and Methods

Membrane embedded alpha helix specific terms were removed from the 2006 Yarov-Yarovoy centroid score function for membrane proteins(96). The modified set of weight terms were used in the stages prior to full atom refinement of a comparative modeling protocol (Fig. S3). The full atom score function used for comparative modeling was *franklin2019*, part of the RosettaMP framework that has been validated for changes in Gibbs free energy given a mutation (DDGmut) and native structure identification of TMBBs (2).

The *franklin2019* full atom core function was used for all trajectories. To account for the acidic environment on the exterior of the cell, the flags “-pH_mode” and “-value_pH 2” from Rosetta-pH were used to calculate the pKa values assuming media of pH 2 and set protonation states of residues outside the membrane (43). The score terms *fa_elec* and *e_pH* was set to 1.0 and added to the franklin2019 score function.

Spanfiles to determine the transmembrane regimes were from Boctopus, a server that uses support vector machines (SVMs) and Hidden Markov Models (HMMs) to calculate TMBB topologies (31). Constraints based on evolutionary similarity were generated from GREMLIN (Fig. S12). Threaded models were based on GREMLIN calculations of evolutionary similarity (71). These weights files were then used in the 2018 RosettaCM protocol (13).

The same protocol was used for comparative modeling of Cyc2 (UniPROT: B7JAQ7_ACIF2) from *Acidithiobacillus ferrooxidans* (strain ATCC 23270 / DSM 14882 / CIP 104768 / NCIMB 8455). Signal peptides in the sequence were detected by SignalP 5.0, a neural network signal peptide detection server (5). Threaded models were generated from 16-stranded TMBBs with known structures identified using GREMLIN as having evolutionarily conserved and coevolved regions with above 95% probability of residue-residue contacts (9; 40; 71) (Fig. S12). Outer membrane porin proteins OprP and OprO from the gram-negative bacterium *Pseudomonas aeruginosa* (respective PDB codes 2ov4 and 4rjw) were used as homologs (65; 80). OprP and OprO are trimeric porins, but earlier molecular weight characterizations of Cyc2 indicated that it is a monomeric protein(6), so one subunit of each of the *Pseudomonas aeruginosa* OMBB proteins was used. These threaded models were hybridized, and the new weight sets were used to *ab initio* fold regions with low structural or sequence homology in a membrane environment. The model used DPLC (1,2-dilauroyl-sn-glycero-3-phosphocholine), a common component of gram-negative bacteria outer membranes with membrane depth of 30 A. The resulting models were then minimized using MPRelax with the *franklin2019* score function. 100 trajectories were run of the CM protocol and the three lowest-scoring models were checked using MolProbity (92). The model with the most favorable protein geometry was selected as the most likely.

The Cyc2 sequence has a known heme c binding domain (74) that was modeled based on homology from a cytochrome c identified by GREMLIN to have the highest coverage and most probable residue-residue interaction (PDB code 2zzs) (14). The cytochrome c554 was from *Vibrio parahaemolyticus,* a bacterium that obtains energy for growth from the oxidation of ammonia to nitrite (14). Heme c was docked via homology and verified by measuring the heme iron-imidazole distances. Both Fe(II) and Fe(III) docking sites were found by homologous templates in the PDB using the Metal Ion-Binding Site Prediction and Docking Server (MIB) (53). Potential metal binding sites were searched in BLAST against proteins with similar functions in other organisms. The ten most similar protein sequences were compared for conserved residues in the potential iron binding sites, and any sites without similar residues conserved across species of iron oxidizing bacteria were discarded (Fig. S8, S9). Amino acid side chains in metal binding sites in conserved regions were repacked in PyRosetta using the flag “in:auto_setup_metals” and the metalbinding_constraint weight set to 1.0 to arrive at more favorable metal chelating geometries as calculated by metalbinding_constraint score term (88) and relaxed using 100 trajectories (Fig. S13). These metal geometries were verified with the CheckMyMetals webserver (100).

Abergel created docked models of the periplasmic proteins in the electron transfer chain downstream from Cyc2. Rcy (PDB code 1rcy) is an oxidizing cupredoxin that likely serves as the immediate downstream partner in the electron transport chain from Cyc2 (87). Cyc1 (PDB code 1h1o) is downstream of Rcy and serves as a “tuner” for mediating electron transfer (1). From experimentally determined interactions, fast Fourier transform (FFT) docking was performed in ClusPro between Rcy and Cyc1 (46; 84) and refined using ROSIE Docking (59) (Fig. S14). Interactions between Rcy and Cyc2 were determined from conserved regions on the Rcy faces known not to interact with Cyc1. Cyc2 was docked onto the likely Rcy-Cyc1 complex (1) using ClusPro for global docking. All residues in the transmembrane domains of Cyc2 were assigned a repulsive score to favor Rcy docking with the cytochrome region. The best-scoring model was processed in ROSIE Docking2 for refinement. Electron transfer across the three docked proteins was calculated using eMap, a server that predicts electron tunneling through electron transfer active (ETA) moieties using graph theory (80).

To model the transient initial binding of Fe(II) on the outermost loops of Cyc2, electron potential was calculated using a Poisson-Boltzmann model and the regions with the most negative potential were identified(8; 24).

## Abbreviations

REU: Rosetta Energy Unit
TMBB: trans-membrane beta barrel protein
Cyc2: gene name for outer membrane cytochrome c protein of *A. ferrooxidans*
Cyc1: gene name for cytochrome c552 protein of *A. ferrooxidans*
Rcy: rusticyanin protein of *A. ferrooxidans*
RMSD: root mean square deviation of atomic positions, given in angstroms (Å)
OPM: Orientations of Proteins in Membranes database (58)
MIB: Metal Ion-Binding Site Prediction and Docking Server (53)

## Acknowledgements

The authors gratefully acknowledge financial support from Rosetta Commons and NSF REU Award 1950697 as well as the U.S. Army Research Office (grant W911NF-18-1-0239).

