## Supplemental Information for "Computational Structure Prediction Provides a Plausible Mechanism for Electron Transfer by the Outer Membrane Protein Cyc2 from *Acidithiobacillus ferrooxidans*"

MVSSSVGFKKKRLIVALAAVGGMALSSGAWALPSFARQTGWSCAACHTSYPQLTPMGRMFKLLGFTTT
NLQRQQKLQAKFGNSVGLLISRVSQFSIFLQASATNVGGGQAVFGPGNSNAGASPNNNVQFPQQVSLFY
AGEITPHIGSFLHLTYSGGGSGAGAGGFSFDDSSIVWTHPWKLGTNNLLVTGVDVNNTPTAMDLWNTTP
DWQAPFFSSDYSSWGHVPQPFIESSAGAGYPLAGVGVYGADIFGPNRANWLYADADVYTNGQGTQVN
PVGGFTAAGPQGRLSGGAPYVRLAYQHDWGDWNWEVGTFGMWSSVYDNTINNTLNKAGGPIDTFDDY
DLDTQLQWLDTNDNNNVTIRAAWVNEQQQFGAGNVISSNSSGNLNFFNINATYWYHDHYGIQGGYRNV
WGSANPGLYGTTYTNSGSPDTSNEWIEASYLPWWNTRFSLRYVVYNKFNGVGSASSNNLGYGASAYNT
LELLAWISY

**Fig. S1: Primary sequence of Cyc2** (UNIPROT: B7JAQ7_ACIF2) (85). Underlined residues have been identified as a signal peptide (5).

atggtgtcat cgtccgttgg ttttaaaaag aaaaggttga tcgtagcatt agcagcagtt 60
ggtggaatgg cgttgtcttc cggtgcctgg gcactgccat cctttgcgcg tcagaccggt 120
tggtcgtgcg ctgcctgtca cacatcctac ccgcagttga cgcccatggg cagaatgttc 180
aaattgctcg ggttcacgac cacaaacctg caacggcaac agaaactcca agccaagttc 240
gggaacagcg tcggtctgct catatcccgc gtatcacaat tttctatttt tctgcaggct 300
tcggcgacca atgttggtgg cggtcaggcg gtgtttggtc ctgggaactc taatgcgggt 360
gcttctccca acaataatgt tcagttcccc caacaggtga gcttgttcta tgccggtgaa 420
atcactccgc atattgggtc gtttctgcat ttgacctact ctggcggcgg cagtggtgcc 480
ggtgccggag gatttagttt tgatgactcc agcattgtct ggacccatcc atggaagttg 540
ggcaccaata atcttttggt cacgggcgta gacgtcaaca atacgccgac tgctatggac 600
ttgtggaata ccacaccgga ttggcaggca ccatttttta gctcagacta ttcgtcttgg 660
ggccacgtac ctcagccatt cattgaaagt tcagcaggtg ctggttaccc attagcgggt 720
gttggtgtct acggagccga tatcttcggg ccaaacaggg caaactggct ctacgcagac 780
gccgatgttt ataccaacgg tcaaggaacc caggtcaacc cggttggcgg ctttactgca 840
gctggccccc agggcagact ttcagggggc gccccttatg ttcgtcttgc ctatcagcac 900
gattggggtg actggaactg ggaggtcggt acctttggca tgtggtccag cgtgtacgat 960
aacaccataa ataacactct caataaagca ggcggcccca ttgatacctt cgatgattat 1020
gatttagata ctcagctcca atggcttgac accaacgaca acaataacgt gacgatccgt 1080
gctgcatggg taaacgagca gcagcaattt ggagcgggga atgtcatatc ttcgaactcc 1140
tccggtaact tgaatttctt caatattaat gccacctatt ggtatcacga ccactacggt 1200
atccagggcg gataccggaa tgtgtgggga tccgccaacc ccggtctcta cggtaccaca 1260
tacaccaata gtggttctcc ggacaccagc aatgaatgga tagaggcttc ctatctgccg 1320
tggtggaata cccgcttctc cttgcgatat gtcgtataca acaagttcaa tggcgttggt 1380
tcggcgtcgt ccaacaacct tggatatggg gcgtctgcgt ataacaccct tgaactgctg 1440
gcctggatat catactag

**Fig. S2: DNA sequence of Cyc2** (ENA: ACK79618) (85)

hbond_sr_bb 1.17
hbond_lr_bb 1.17
rama 0.15
omega 0.2
rg 0.1
vdw 3.0
Menv 2.019
Mpair 1.0
Mcbeta 2.5
cenpack_smooth 1.0
cart_bonded 0.05
atom_pair_constraint 0.5
Mlipo 1.0
rsigma 1.0
sheet 1.0
ss_pair 1.0
hs_pair 1.0

**Fig. S3: Modified RosettaCM stage 2 centroid level weights**. Modified from membrane RosettaCM protocol (13).

| | *Existing* | *New* |
|---|---|---|
| *1bxw* | 18.91 Å | 4.45 Å |
| *1qd6* | 16.41 Å | 6.51 Å |
| *1kmo* | 13.25 Å | 8.37 Å |

**Fig. S4: RMSD for lowest-scoring trajectories for RosettaCM modeling test set**. To benchmark these altered scoring functions, we used the modified TMBB CM protocol to create comparative models for three test TMBB proteins that have experimental structures available in the Protein Data Bank. Test case proteins were chosen based on availability of membrane information in the crystal structure and similar low percentages of sequence homology (less than 20% sequence identity). Homologous proteins, on which the sequence was threaded had below 30% sequence identity. This replicated conditions similar to the available homologs for Cyc2. The C-alpha RMSD values for the lowest scoring models after the low-resolution step using existing membrane scorefunctions (Existing) were compared to our new beta-barrel membrane score functions (New). The new model yield small RMSD values for each protein.

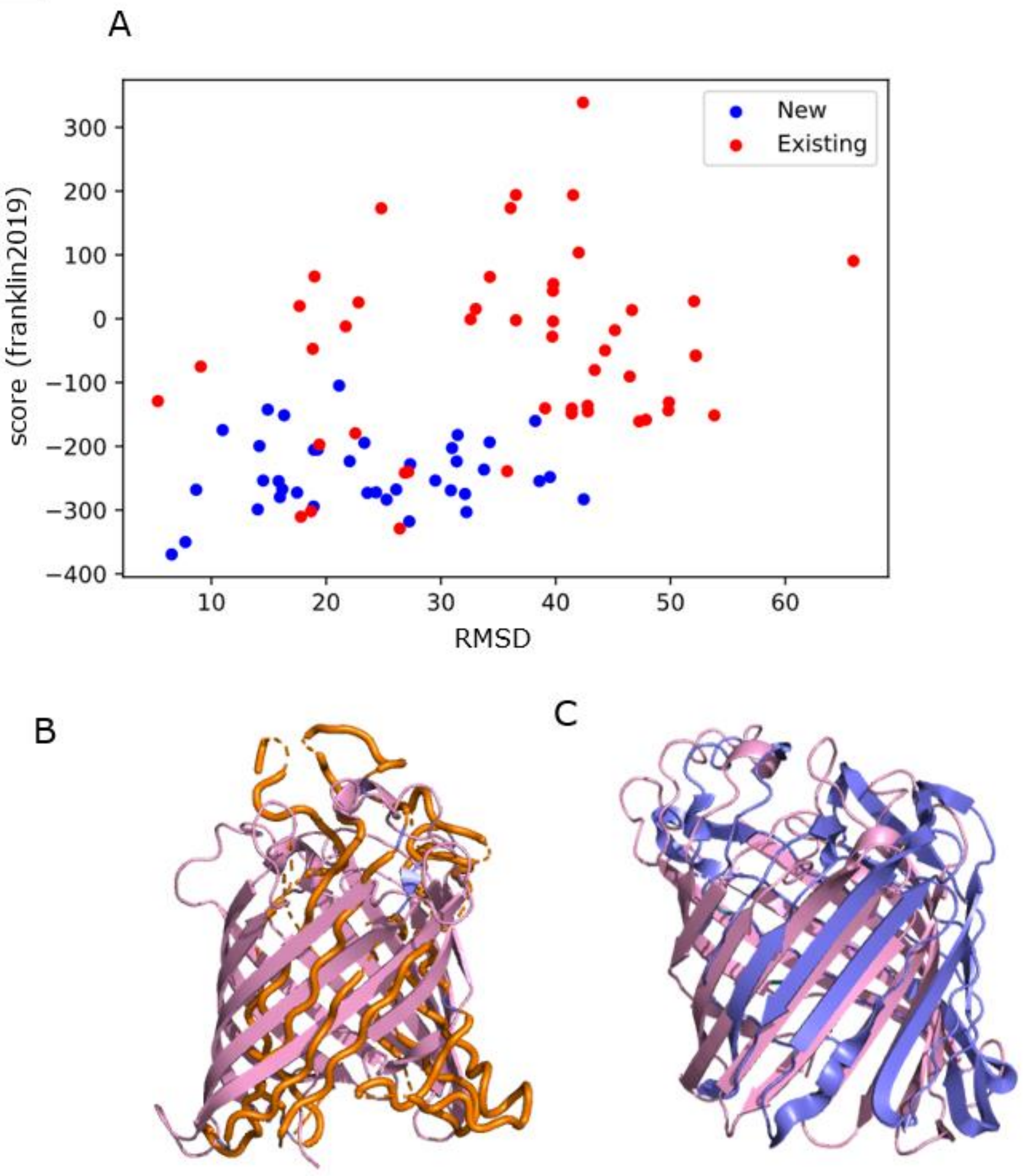


**Fig. S5: Sample benchmark testing on experimentally determined TMBB structure.** Comparisons of RMSD between the new score function and the existing score function for the test set were done in Excel using a one-way ANOVA. Score vs. RMSD plots showed a narrower energy funnel for our TMBB score functions A: Score vs. RMSD plot for TMBB 1qd6 using existing centroid-level score function and modified centroid-level score function for the intermediate folding steps in RosettaCM. Final models were scored with the franklin2019 score function. B: Comparison between experimental (pink) and lowest-scoring CM 1qd6 trajectory with original scoring protocol (orange), with RMSD = 16.41. C: Comparison between experimental (pink) and CM 1qd6 with new scoring protocol (purple), with RMSD = 6.51.

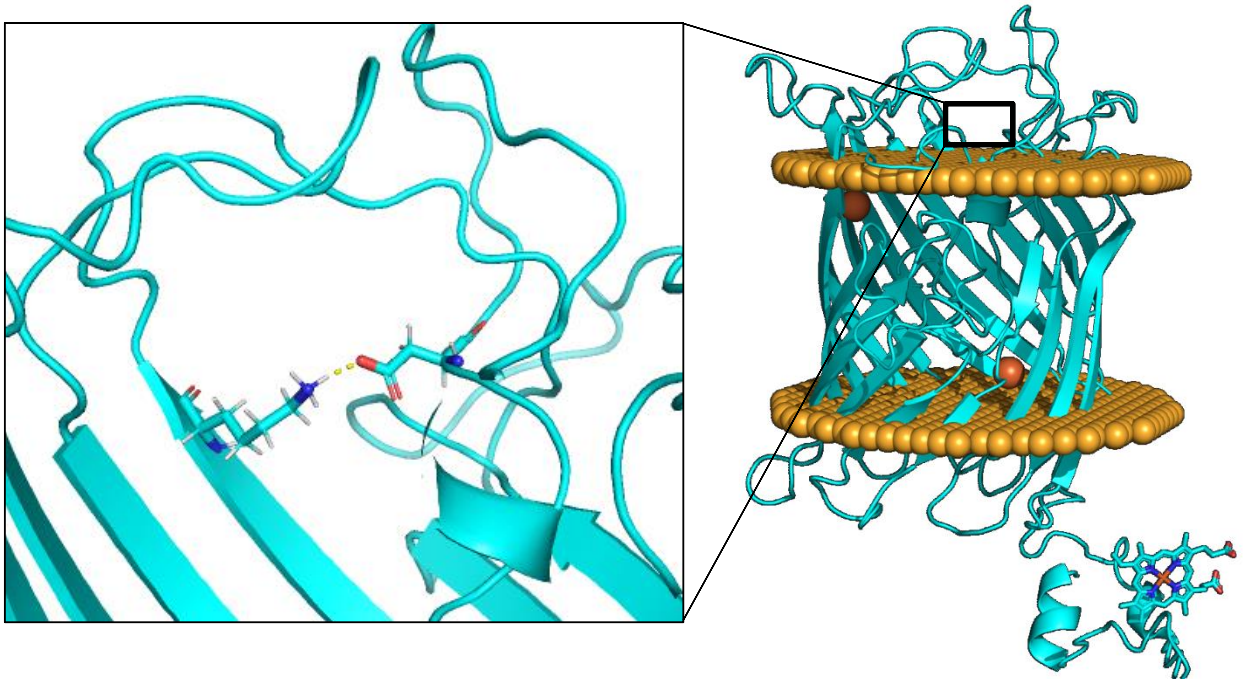

**Fig. S6**. **Effect of pH on Modeling of Cyc2.** Modeling at pH 2 did not modify the backbone structure, however the low pH environment eliminated the formation of a salt bridge predicted in the structure at pH 7. At pH 7 a salt bridge could form between NZ of LYS424 and OD2 of ASP185.

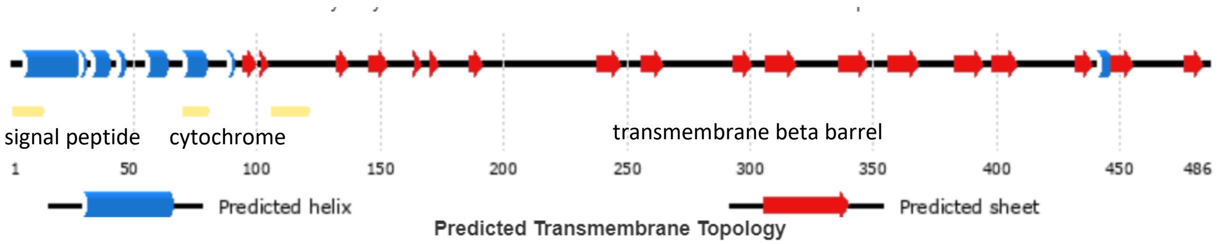


**Fig. S7: Domains of Cyc2 sequence.** Computed by PSIPRED secondary structure prediction. Cyc2 has an anomalously high molar extinction coefficient (144520.00 $M^{-1}cm^{-1}$), indicating many aromatic residues. These residues can accept and donate electrons, indicating that Cyc2 is a likely electron transport protein.

| NCBI accession number | Species |
|---|---|
| ACK79618.1 (TARGET) | *Acidithiobacillus ferrooxidans* |
| WP_114282823.1 | *Acidiferrobacter thiooxydans* |
| WP_070079636.1 | *Acidihalobacter prosperous* |
| OYV75648.1 | *Chromatiales bacterium* |
| HAH22798.1 | *Prolixibacteraceae bacterium* |
| PYO94379.1 | *Gemmatimonadetes bacterium* |
| TSA29379.1 | *Ignavibacteriales bacterium* |
| NJD23605.1 | *Melioribacter sp.* |
| ODU99277.1 | *Rubrivivax sp.* |
| TLZ45134.1 | *Gammaproteobacteria bacterium* |
| AOY94926.1 | *Cupriavidus sp.* |

```
cyc2                ---------------------------------------LPSFARQTGWSCAACHTSYPQ       21
WP_114282823.1      MFHREKNSMAFKIPRLEPRKVLALTSIIGGLAMSANAWALPSFARQTGWSCATCHTSFPQ       60
WP_070079636.1      --------MSISIHRH---LRL-PAGAISLLLPFAPAHAVPSFARQLGVSCAACHTAYPQ       48
OYV75648.1          ---------------------------MALGLASTAVYAVPSFARQTGMPCSACHTVFPE       33
HAH22798.1          --------------GI---NILKLFILMIILAQAGVSKATPSFARQTGMSCTACHYSFPE       43
PYO94379.1          -----------------------MRAYDAIYAVVRTKLGIPSFSRQTGLACNVCHTAFPM       37
TSA29379.1          --------MGSKKATG---IIRLLFLITVLLIFSNQTKAIPSFARQTGMSCNACHTIFPE       49
NJD23605.1          --------MDSKKNTI---VNSLLFLLTVFLFFSNQTQAIPSFARQTGMSCNTCHTIFPE       49
ODU99277.1          -------------MAT---LGM-AGGAL--LLAPGSAQAVPSFARQTGMACEACHTVFPE       41
TLZ45134.1          --------MKSGAGNT---RAL-CGAALLSVLTAPAALAVPSFARQTGMACEACHTVYPE       48
AOY94926.1          --------MKARARYW---IAFGAAFALLIAALPQSGMAVPAFARQTGMACVACHVNFPE       49
                                                            *:*:** *  * .**  :*

cyc2                LTPMGRMFKLLGFTTTNLQRQQKLQAKFGNSVGLLISRVSQFSIFLQASATNVGGGQAVF       81
WP_114282823.1      LTPMGRIFKLMGYTTTNLQPQQKVEAKVGNVVHLLLPRISQFSIFVQASDTHVAGSQQAL       120
WP_070079636.1      LTPFGRQFKLLGYTMQNEPTVKADNGK-----RLDIDRWAPLSLMVMVSDSLPQHPGPGN       103
OYV75648.1          LTAFGRSFKLNGYTLTGMKQIESTGTS-G---NVRINAIPPLSAMLQTGFTHLNKAVPGQ       89
HAH22798.1          LTPFGRQFKLNAYTMTMMNTIESKQDS-DKVTRLKLLSYLPLSAMVQTSFSSNAKAVEGT       102
PYO94379.1          LTAFGRQFKLNAYTLTGLQTIGPTET--T-PSPLKINLIPPVSTMLQTSFTQTSKAQPGT       94
TSA29379.1          LNAFGRQFKLNGFTLVATEIIQAQADS-E-TTILSLPKSSPFSAMLQASYTYKSKEQPAT       107
NJD23605.1          LNAFGRQFKLNGFTLVGMETIKAMSDS-E-TTILSLPKLSPFSAMLQASYTYKSKEQPGT       107
ODU99277.1          LTPFGRRFKLNGYTIDNLPQVSGMNAN--KDQTLALNQLPPLSFMFQTSYTKTKTAVPDS       99
TLZ45134.1          LTHFGRVFKANGYVLANLKQVRDVTAK--KEQLLELGQTPPLSIMVQASYTQLSTTVPDL       106
AOY94926.1          LTPFGRFFKLTGYTLSNNR-------------------TIPLSAMVQVSRTSSRTVDQA-       89
                    *. :** **  .:.                           .* :. .. :

cyc2                G-----PGNSN-----AGASPNNNVQFPQQVSLFYAGEITPHIGSFLHLTYSGGGSGAGA       131
WP_114282823.1      G-----APART-AKGGQPVGTNNNLEVPQQVSLFYAGEVTPHVGDFLHITYNGQ-----S       169
WP_070079636.1      ---------------------DASQVEFPNQLSLFYAGAITDHIGTFLQYTA-GD-----G       137
OYV75648.1          ---------------------QNDNVEFPQALSFYFAGEISPHMGSFLQVTYTQP-----D       124
HAH22798.1          ---------------------QNNSIAFPQQISMFYAGQITPHIGSFIQMTYDGQ------       136
PYO94379.1          ---------------------QNGNVEFPQELSVFFGEAISPRLGTFIQITYDGA-----E       129
TSA29379.1          ---------------------QNGNFSLPQQLSFFIAGALTPKVGGFIQATYSDQ-----D       142
NJD23605.1          ---------------------QNGNISFPQQLSFFVAGALTPKVGSFIQVTYSDQ-----D       142
ODU99277.1          QAGVIPPGGTAAPVSADALAKNGQVLFPQQASLFYAGRIAPNFGAFVQMTYHGT-----A       154
TLZ45134.1          SQ------------SAPGVAQNGTAGFPQQLSLFYAGKIAPHFGAFVQLTYAND-----S       149
AOY94926.1          N---------------FDF-VRNDDLALQQASVFLAGRIFDHVGTFTQWTYDGI-----A       128
                                          .     :  *.: .  :  ..* * : *

cyc2                GGFSFDDSSIVWTHPWKLGTNNLLVTGVDVNNTPTAMDLWNTTPDWQAPFFSSDYSSWGH       191
WP_114282823.1      GTFAFDDSSIVRTQAWKLGLHNTLITGVDVNNTPSATDLWNTSPDWQAPFFTSNYTAEGA       229
WP_070079636.1      GAFGLDNTDVRYANQTSLG-STSVIYGIDLNNNPTVQDVWNSTPAWQFPYFTPGDTFTTA       196
OYV75648.1          DHFGFDMADFRYANRTTLG-ERGLVYGVTLNNAPGMEDVWNTTPMWTYPYTASDTA----       179
HAH22798.1          -VFGMDNAEIRYANRTSLG-STSLLYGVTLNNNPTVQDVWNTTPAWGFPTASSDAA----       190
PYO94379.1          GTLGVDNIDLRYANHAKVL-SKRTIYGITVNNSPTVQDVWNSTPVWGFPFGSSGVA----       184
TSA29379.1          GAFKLDNAEIRYADQTELA-SESFTYGVTLNNNPTVQDLWNTVPAWRFPYAGSSVS----       197
NJD23605.1          GSFGIDNAEIRYADQTELA-SESFTYGITLNNNPSVQDLWNSTPAWRFPYAGSSVS----       197
ODU99277.1          GTFGWDNTDIRYARAV----NDKFLWGLSFNNNPTVQDLWNTTPAWQSPFDQRSATA---       207
TLZ45134.1          GTISIDNTDLRFADMMVLPSEQSLVYGISLNNNPTVQDLWNSTPAFGFPYASSNAV----       205
AOY94926.1          HRAALDNTDIRAAWHL-AENDIDFIYGVTVNNNPTVSDVWNSTPAFGFPFASSSVS----       183
                         *  ..  :             *: .** *   *:**: * :  *    .

        * :
```

```
cyc2                VPQP----FIESSAGAGYPLAGVGVYGADIFGPNRANWLYADADVYTNGQGTQVNPVG-G      246
WP_114282823.1      VPTT----FIGSSPGAAFPLIGLGAYAADIVGPNRANWFYAEADAYNNSEGTGAAPNQGG      285
WP_070079636.1      GGAAVPATLIEGA-LA-QQVAGLTAYGY------FDNTYYVEAGVYHGMNNP--------      240
OYV75648.1          -PTPAAGPLVN---MF-MNVAGLGGYAL------WDNHWYGAATVYRSAPLG-------Q      221
HAH22798.1          -NSPAKSTLIE-N-LG-MQVAGLGAYAL------IDNLLYGELTLYRSSPQG-------A      233
PYO94379.1          -PTPSGSPLINEG-LA-QSVAGIGTYAL------WDDHLYGEVSVYRSSPQG-------G      228
TSA29379.1          -PTPSASPLIDGG-LA-QNVAGLGVFSM------WNNLLYTEISVYKSAQQG-------S      241
NJD23605.1          -PEPSASPLIDGG-LA-QNVAGIGVFSM------WNNLLYTEISVYKSAQQG-------S      241
ODU99277.1          -PGPSATTMIDGG-LAGAGVAGLSIYGS------WNDSIYVELGAYRSAPA---------      250
TLZ45134.1          -VSPLAGTEIDGG-LA-QDVAGLSGYLF------WNESLYAEVGGYRSAKQGAANALTGA      256
AOY94926.1          -VTPAASTLIDGG-LA-QQVAGLAAYAF------WQRSVYAEFGFYRTADGALSVFRTGQ      234
                             :         : *:  :             *     *

cyc2                FTAAG---PQGRLSGGAPYVRLAYQHDWG-DWNWEVGTFGMWSSVYDNTINNTLNKAGGP      302
WP_114282823.1      FFEGGSNSLFGQLAGTAPYLRLAYQHDWS-DWNWEVGTYDMWSRVYASPISNA-------      337
WP_070079636.1      ST-DSPANGGQYIKGMAPYWRLAYTGQAG-NSNWEVGTLGLIAD-------VPVDGPTGP      291
OYV75648.1          GA-AP---SAGSVRNVAPYWRFAWQGYLPNRAYLEVGTYGLYADF-PRG--MMGMGTPGL      274
HAH22798.1          AN-PADSTSVMTVRGVAPYWRVALQHAWG-NNYAELGTFGIASN-------NFVNGISGP      284
PYO94379.1          PH-PPDATSTGIMKGVTPYWRLAYQRTFG-TQYIELGTFGMASQ-------LYPTGVTGP      279
TSA29379.1          HN-PPDSTSAGVLKSLAPYWRIALQKQFG-DCYLELGTFGLSAT-------MFPAGITGL      292
NJD23605.1          PN-PPDSTSSGILKSLAPYWRVALQKQFE-DFYLELGTFGLSAT-------MYPAGVTGL      292
ODU99277.1          AA-QIDSTSANVVSGMAPYWRLAYERQWG-RSSWEAGLYGIDAKLYPGG----GNALSGP      304
TLZ45134.1          AG-PLDGTASNVIEGVSPYWRVAYEYNHE-RHSIEAGLYGADFKLLPGA--STGTPLRGP      312
AOY94926.1          DI-NT-PGGVARLSGASPYWRLAYNHEWG-ANSLMLGTFGMIADRYPDN--T---LPGTP      286
                                : . :** *.*             *  .

cyc2                IDTFDDYDLDTQLQWLDTN-DNNNVTIRAAWVNEQQQFGAGNV--------ISSNSSGNL      353
WP_114282823.1      VNRFYDYDLDSQLQWLDIN-DNDNVTVRANLIHEDANFAAGAL--------NTSLTHGQL      388
WP_070079636.1      TDKFTDIGFDGQYQWLAGK---NTVTVHGAYYHEHQNLNGTTALTGT------SYGSQHL      342
OYV75648.1          TDKYIDLAVDTSYQQPLS--GTHLLSLHGVYIHENRTLDSSFAKG------LSSNRTDTM      326
HAH22798.1          IDKFTDVGFDLQYEHTLNT---GTLVLHSSLIRETEKRNSTIN-------------NFHF      328
PYO94379.1          TDHFSDVGADLQYERHAGKNGAGTFVVHASYIDERQTLDASFGGG------ASANAKNTL      333
TSA29379.1          TDQYSDVGFDLNFEKAFGA---DMLSARGSWIHESRTLDASVANG------SAFNTSGNL      343
NJD23605.1          TDQYSDVGFDLNFEKAFGT---DMLSARGSLIHESRTLDASVANG------EAFTTSGNL      343
ODU99277.1          TNHYRDWAIDSQYQYIGDE---HMISVLATYISEKQTLDASYGVL----G-TAANPTNDL      356
TLZ45134.1          FNRFKDVAEDIQYQFIADE---HQVTVAGTRIHENMSLDASFAAT----PAASANPKDDL      365
AOY94926.1          TDRFSDYALDAQYQYLTLP---HAFTAQAAWIYEKQNWRASFPSGGIGAGPTPANPTDHL      343
                     : : *   * . :          .   .    *     .                   :

cyc2                NFFNINATYWYHDHYGIQGGYRNVWGSANPGLYGTT---Y---------------TNSGS      395
WP_114282823.1      NTFNLNATYWYHDEYGAQGGFQDVTGTANSSFWGGN---V-------------YTSANGS      432
WP_070079636.1      DTYNVSATYYYRRMYGATLAYLGASSSSNAVDQG----SLAGPAGASTASGIPTTYAPGD      398
OYV75648.1          QQVRVDGNYEFSHHAQVSLGYFNTWGSTDAAFYATTPGAVDN-------------SSSGN      373
HAH22798.1          NSFKIDGNLYLKNGLGATMGYFNSSGTKDAD--------VVE-------------SSTNK      367
PYO94379.1          NTVRADAAWLTPTRWGGSVGVFNTSGTADTLLYA--PGAVTG-------------NATGK      378
TSA29379.1          NSFRVVGNYYLHSQIGFSLGYFSMTGDGDAILYA--PTSVSG-------------SANGL      388
NJD23605.1          NSLRFVANYYMHSQIGFSLGYFSMTGDGDNILFA--PTSVSG-------------SANGL      388
ODU99277.1          KTARIGVNYYYHRRYGGALGYFSTTGSADSGLYA--PAPFTG-------------FANNK      401
TLZ45134.1          TTTRLWATYYYRRKIGGTLGYFSTTGSGDAVLYP--PNAAGGPGVV--------TSANGS      415
AOY94926.1          TTFKARASYMYQRKYGGTLAYFSTTGNADPGLYA--PAPVTG-------------SANGY      388
```

```
cyc2                PDTSNEWIEASYLPWWNTRFSLRYVVYNKFNGVGSAS-----SNNLGYGASAYNTLELLA      450
WP_114282823.1      PNTTDEWVEASYLPWWNTRLSVRYTVFNKFRGLTG---------NNGVSPSKFNTIELLA      483
WP_070079636.1      RGASAYIAQLDYVPWYNTQFSLQYMAFNKVNGTTT-------------DAALNNYFMLGM      445
OYV75648.1          PGSAGLITEVDYLPWDNTKFSLQYTAYTKFNGAGSNYNGA------GRSASDNNTLYLNS      427
HAH22798.1          PNSNGFIYQIEYLPWYNTKFSIQYITYSMFDGSSTNYNGA------GRNASDNNTLYLLA      421
PYO94379.1          PNSNGVIAELQFMPWINTRFSLQYVAYQKFNGGTSDYDGA------GRSASANNTVYALV      432
TSA29379.1          PDSGGFTAEFDYLPWLNTKLSVQYVAYNKFNGGSNNYDGE------GRNASDNNTLYLLC      442
NJD23605.1          PDSNGFTAEFDYLPWLNTKLAIQYVAYNKFNGGDNNYDGQ------GRNASDNNTLYLLC      442
ODU99277.1          PDSKGWTAELDWVPYENTKFALQYTNYSKFNGGSTNYDGTATPDHPGRNASDNNTLYLLG      461
TLZ45134.1          PDTRGWIAEVNYLPWLNTKLTAQYVRYNKFNGASSNYDGA------GRDASDNNAWYLLL      469
AOY94926.1          PDSRGLIFELDYLPHPQVKLALQYTWFLKFNGAHANYDGN------GRNAQDNNTLYLLA      442
                     .:     : .::*  :.::: :*  :  . *                .    *

cyc2                WISY    454
WP_114282823.1      WIAY    487
WP_070079636.1      WYAY    449
OYV75648.1          WLMW    431
HAH22798.1          WFNF    425
PYO94379.1          WLMF    436
TSA29379.1          WVAF    446
NJD23605.1          WVAF    446
ODU99277.1          WINF    465
TLZ45134.1          WFAY    473
AOY94926.1          WFAF    446
                    *  :
```

**Fig. S8: CLUSTAL O multiple sequence alignment.** Measured across top ten most closely related species cytochromes to Cyc2.

| *Rank* | *Binding Residues* | *Template* | *Score* | *Conserved* | *Ligand* |
|---|---|---|---|---|---|
| *1* | 119H , 137D , 138D | 1w69A1 | 1.175 | TRUE | Fe(II/III) |
| *2* | 367H , 368Y | 2pt2A1 | 0.906 | False | |
| *3* | 308D , 335E , 336Q | 4b2oA1 | 0.858 | TRUE | Fe(II/III) |
| *4* | 269H , 277E | 4hr4A1 | 0.748 | False | |
| *5* | 12C , 15C | 1h79A1 | 0.744 | TRUE | Heme |
| *6* | 43Q , 366D , 367H | 3ovpA1 | 0.711 | False | |
| *7* | 365H , 368Y | 1ey2A1 | 0.708 | False | |
| *8* | 229D , 231D | 3ak9A1 | 0.703 | False | |
| *9* | 324N , 365H | 2bq8X2 | 0.691 | False | |
| *10* | 267Y , 269H | 2xgfA1 | 0.633 | False | |

**Fig. S9: Possible metal binding sites.** Top MIB ranked binding site predictions, conserved across related cytochrome c membrane proteins, and possible binding ligands. Cut offs for conservation were E-value < 3e-52 (top ten species identified by BLAST).

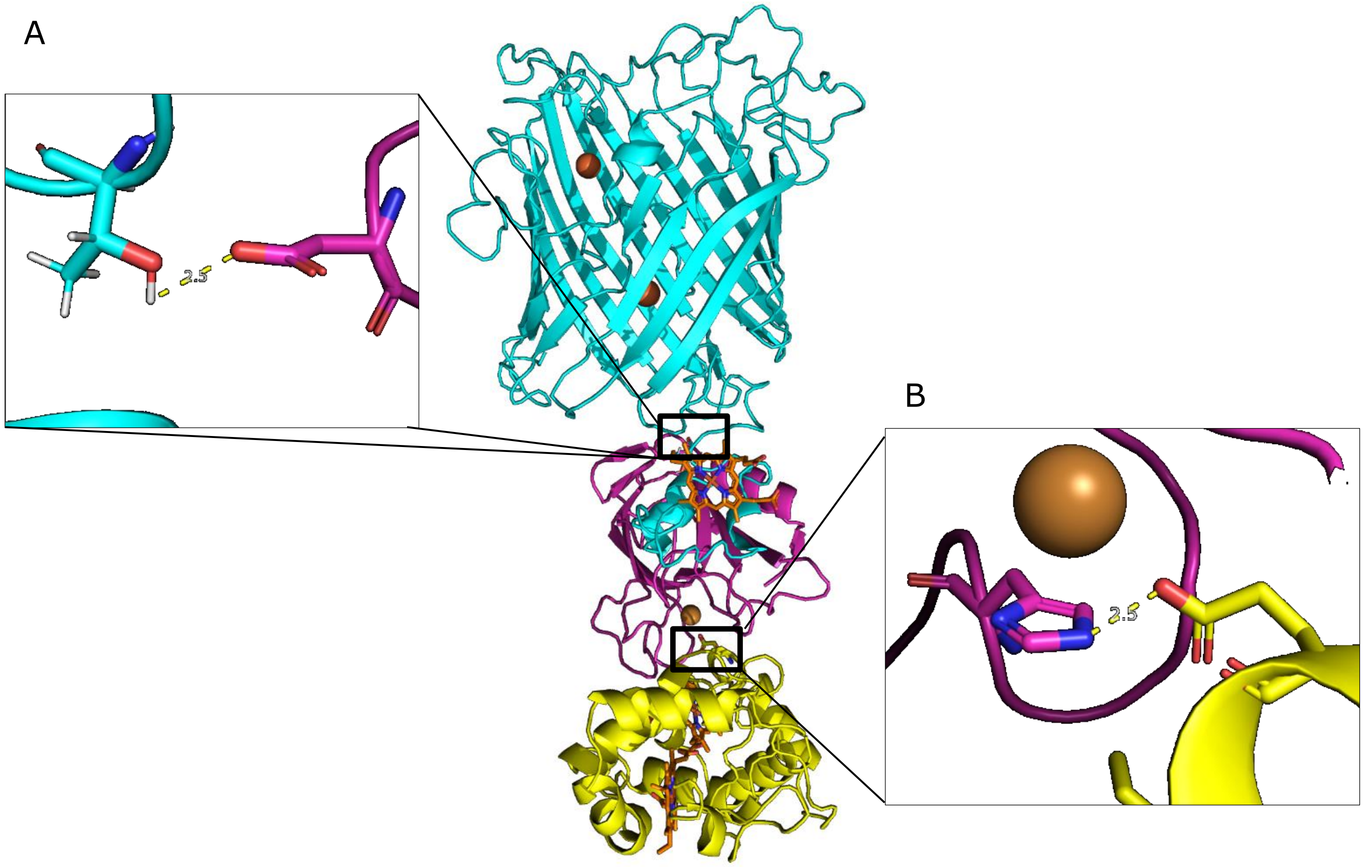


**Fig. S10**. **Identification of possible hydrogen bonding sites between docking partners in Cyc2 electron transfer chain.** A: The OD2 atom of D58 on Rcy was found to form a hydrogen bond with the proton on OG1 of T36 on Cyc2 (distance 2.5 A). B: Hydrogen bond formation is plausible between the NE proton of rusticyanin H143 (carbon backbone in magenta) and the OE proton of Cyc1 E121 (carbon backbone in yellow), creating a docking interface for electron transfer from the bound copper ion in rusticyanin.

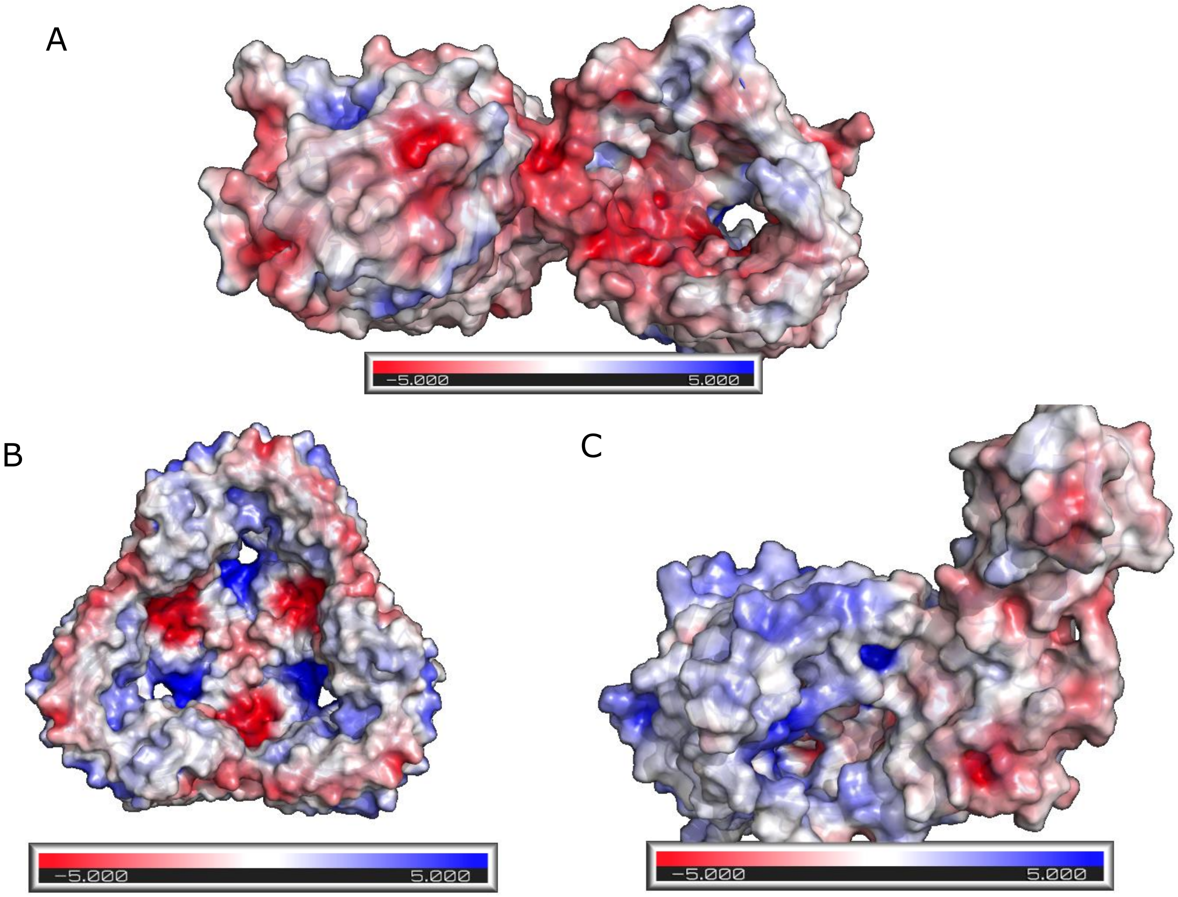


**Fig. S11: Poisson-Boltzmann potential calculated for anion-specific, cation-specific, and nonspecific TMBB porins.** Poisson-Boltzmann potential diagrams created using PDB2PQR, where blue indicates regions of positive potential (> +5 kT/e) and red depicts negative potential (< −5 kT/e). (24). A: Cation selective pathway of OmpF porin (PDB ID: 3HWB). Pore facing residues have high negative potential, shown in red (21). B: Anion selective pathway of Omp32 porin (PDB ID: 2FGQ). Pore facing residues have high positive potential, shown in blue (98). C: Filamentous hemagglutinin transporter protein (PDB ID: 3NJT). There is no strong Poisson-Boltzmann potential gradient within the protein (20).

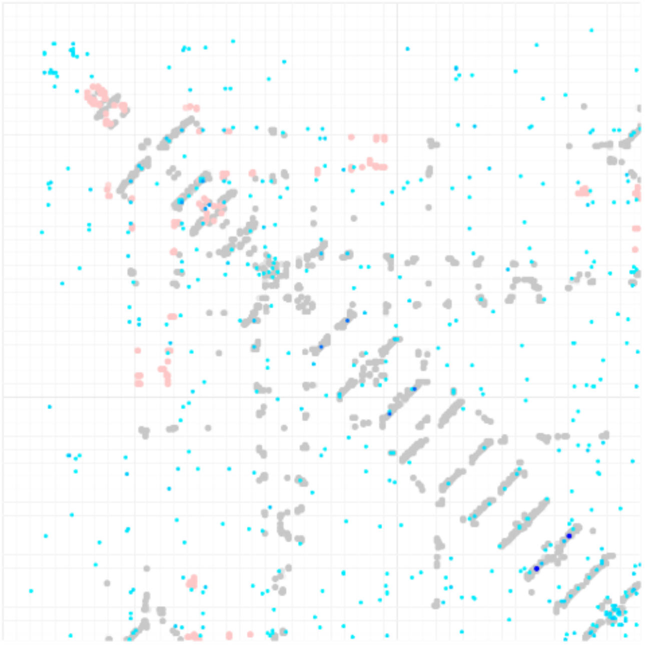

**Fig. S12: Predicted residue-residue contacts between Cyc2 sequence and 2o4v.** Computed by GREMLIN to likely homolog 2o4v, as calculated by evolutionary similarity.

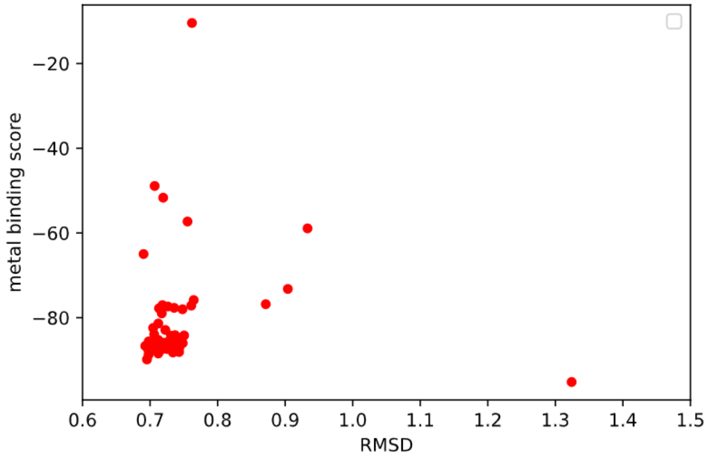


**Fig. S13**. **Metal-binding score vs RMSD.** This plot shows 100 trajectories of FastRelax on residues homologous to metal chelating residues.

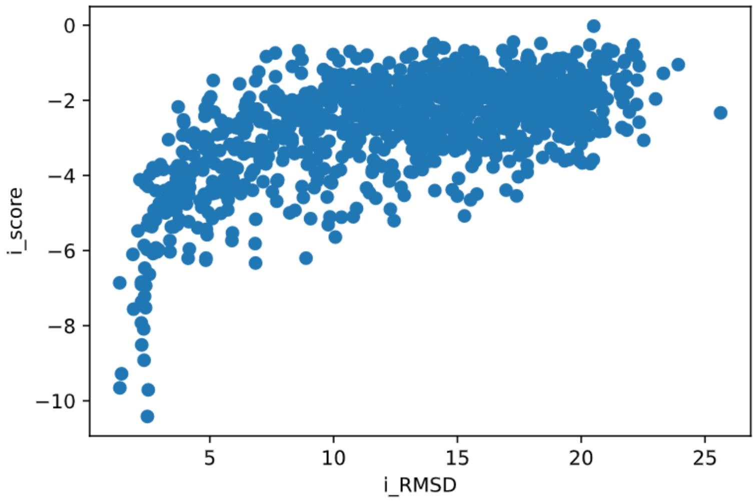


**Fig. S14**: **Funnel plot of interface score vs. interface RMSD for docking Cyc2 model against known Rcy-Cyc1 complex.** Calculated for docking Cyc2 model against Rcy-Cyc1 complex, which has been previously documented (1)
